# A sexual dimorphism in the spatial vision of band-winged grasshoppers

**DOI:** 10.1101/2020.09.18.303784

**Authors:** Alex B. Duncan, Brae A. Salazar, Sara R. Garcia, Nicholas C. Brandley

## Abstract

Visual acuity (VA) --- a measurement of the fineness or coarseness of vision --- correlates with the size of an animal, with larger species often possessing sharper vision. However, it is unknown whether the same relationship between visual acuity and size holds within a species when individuals differ consistently and substantially in size, such as through a sexual size dimorphism. Here we examine the visual acuity of three species of sexually dimorphic band-winged grasshoppers, in which females are the larger sex (*Arphia pseudonietana, Dissosteira carolina*, and *Spharagemon equale;* total n = 98). Using a radius of curvature estimation method, we find that females have ∼21% finer vision in the most acute region and axis of the eye than do males. Further explorations of the eyes of the species showing the greatest size dimorphism (*D. carolina*) suggest that this VA dimorphism is driven by females having larger eyes with more ommatidia. In contrast to many flying insects where males have finer vision to acquire mates, our study is one of the first to demonstrate a female-biased sexual dimorphism in acuity. Given the number of species in which females are larger than males, our results suggest that differences in VA between the sexes may be more common than currently appreciated.

## 1. INTRODUCTION

Animal behavior is driven by what information they can perceive, which itself is limited by their sensory systems (Partan and Marler, 2002; Uexküll, 2013). Therefore, studying how an animal responds to stimuli requires understanding what information it can perceive (Caves et al., 2019; Jordan and Ryan, 2015). Notably, animals differ not just in the presence or absence of senses, but in the fineness with which they can parse information within a sensory modality. For example visual acuity (VA) --- defined as the ability to perceive static spatial detail and used as a measurement of the fineness or coarseness of vision --- varies by up to four orders of magnitude between species (Caves et al., 2018; Land and Nilsson, 2012), making it a promising parameter for further exploration of within sense variation.

Between species, VA changes with both the ecology and morphology of a given species. Ecologically, VA is finer in predatory animals (Veilleux and Kirk, 2014) and those in high-photon environments (Caves et al., 2017; Veilleux and Kirk, 2014). Morphologically, larger eyes (and through correlations between eye and body size, larger bodies) are typically associated with finer vision regardless of taxonomic level, with studies supporting this at the kingdom (Caves et al., 2018; Land and Nilsson, 2012), superclass/class (Caves et al., 2017; Kiltie, 2000; Veilleux and Kirk, 2014), order (Rutowski et al., 2009), and family (Jander and Jander, 2002) levels.

Despite numerous examples of VA varying with size between species, surprisingly fewer studies have examined whether the same principles hold within a species (although see Corral-López et al., 2017; Spaethe and Chittka, 2003). VA --- like other aspects of vision --- can vary within a species in other contexts. For example, numerous within-species VA studies have focused on mate pursuit in flying insects. In these cases, males have finer vision in specialized regions designed to spot potential mates (Kirschfeld and Wenk, 1976; Land and Eckert, 1985), which is accompanied by neurological and behavioral specializations. However, we know little about within-species variation in VA outside of this context. Another way VA may vary within a species is through a sexual size dimorphism. Size dimorphisms may have large effects on vision in small animals with compound type eyes (such as insects). Unlike camera eyes that have a single larger lens, compound eyes have numerous smaller lenses for each of their ommatidia. As a result, the small overall size of compound eyes and their numerous lenses may leave them diffraction-limited (Barlow, 1952; Kirschfeld, 1976; Land, 1997). Thus increases in eye size often improve optical performance (Jander and Jander, 2002; Rutowski et al., 2009).

Based on both 1) the interspecific correlation between size and VA and 2) the optical limitations outlined above, one could predict that sexual size dimorphisms may be accompanied by the larger sex having finer vision. However, this may not always be the case, as larger eyes can optimize other visual parameters (such as sensitivity) instead of VA. Additionally, eye size may not scale with body size within a species. An ideal animal group to examine whether size dimorphisms may be accompanied by VA dimorphisms are the band-winged grasshoppers. Band-winged grasshoppers (subfamily Oedipodinae) are a morphologically diverse subfamily of ∼200 diurnal species known for their colorful hindwing patterns (Otte, 1970; Otte, 1984). Many species of band-winged grasshoppers show a size dimorphism; however, little is known about their vision. Here, we first examine VA in the most acute region of the eye across three species of band-winged grasshopper--- *Arphia pseudonietana* (Thomas, 1870), *Dissosteira carolina* (Linnaeus, 1758) and *Spharagemon equale* (Say, 1825) --- to determine if the larger females have finer vision. In a second experiment, we then further examine eye and receptor scaling in the species that showed the greatest size dimorphism (*D. carolina*) to better understand whole eye changes between the sexes.

## 2. MATERIALS & METHODS

### 2.1 Experiment I: VA variation by biological sex across three species

#### 2.1.1 Study Organisms

To examine how biological sex influences visual acuity in band-winged grasshoppers, individuals of three species were collected during the summer and early fall (June -October) of 2016 and 2017 in Colorado Springs, CO. *Dissosteira carolina* (n = 16 males and 16 females) were collected from an urban site near manicured grass, while *Arphia pseudonietana* (n = 18 males and 18 females), and *Spharagemon equale* (n = 15 males and 15 females) were collected at a high-altitude grassland site. Grasshoppers were euthanized using ethyl acetate and then stored at ∼ -20°C prior to imaging. Total length (head to the tip of the forewings; *mm*), weight (g), and biological sex were recorded for each individual.

#### 2.1.2 Estimating Visual Acuity

We measured visual acuity (VA) of all three species via a modified radius of curvature estimation (Bergman and Rutowski, 2016). Briefly, this method estimates VA by calculating how many ommatidia view a given angle of visual space.

To calculate VA, eyes were imaged at 40X magnification using AmScopeX for Mac MU (MW Series 05/26/2016; United Scope LLC; Irvine, CA, USA) under diffuse lighting conditions (LED312DS; Fotodiox Inc. USA; Gurnee, IL, USA) with a M28Z zoom stereo binocular microscope (Swift Optical Instruments Inc.; Carlsbad, CA, USA) and AmScope 14MP USB3.0 digital camera (United Scope LLC; Irvine, CA, USA). Because grasshopper eyes are not spherical and therefore have different curvatures in each axis, we used a lateral image to measure curvature ---and ultimately VA--- in the axis perpendicular to the horizon (y axis), a dorsal view for the axis parallel to the horizon (x axis), and one anterior view of the flattest region of the eye to measure facet diameter (Figure 1). For consistent positioning, physical attributes were used for orientation in each image (lateral image = the inside eye edge, dorsal image = the top of the eye, anterior image = the center of the eye).

**Figure 1:**
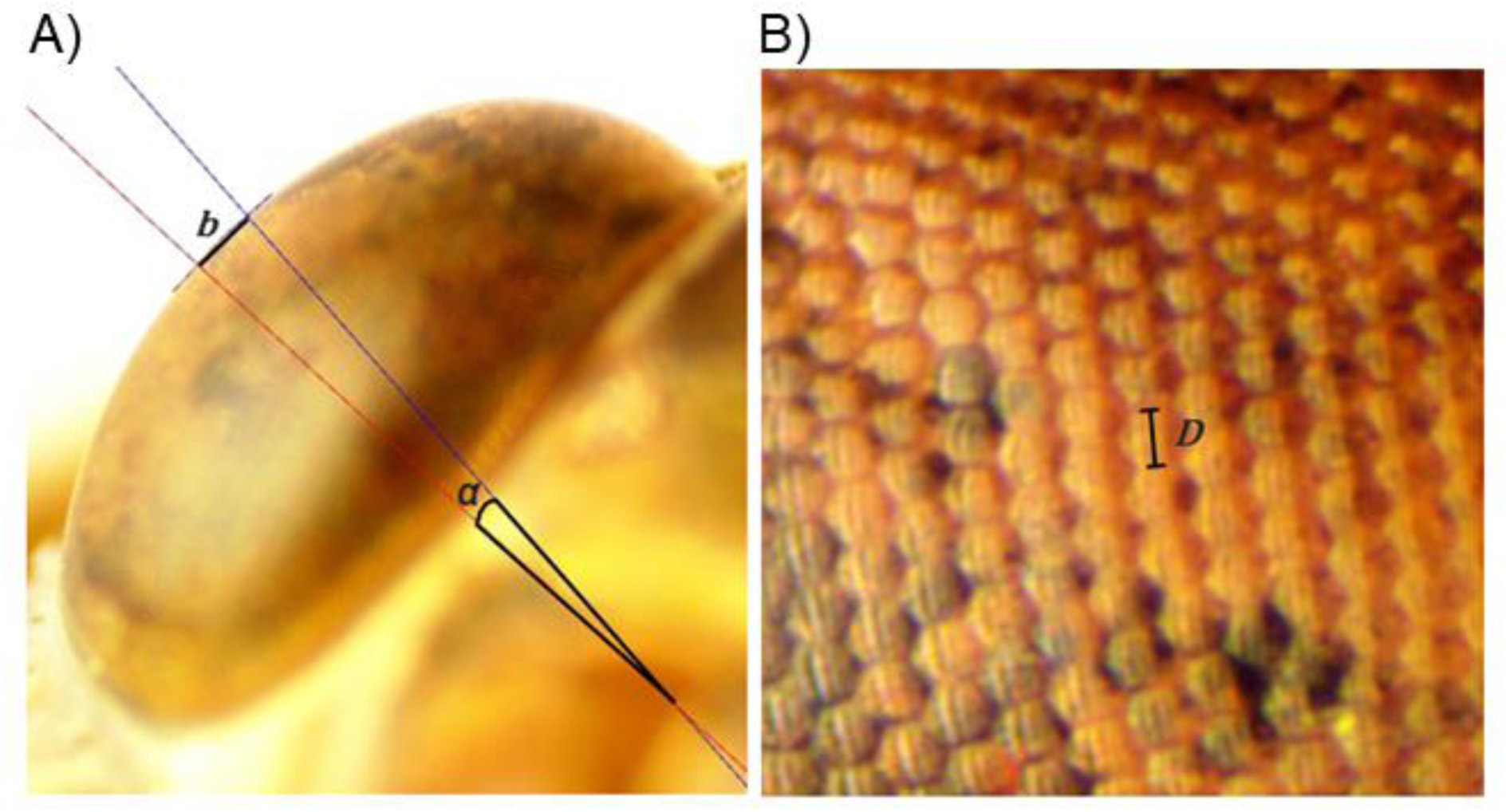
An example of the radius of curvature estimation used to estimate visual acuity in band-winged grasshoppers. A lateral view of the eye with the edge in focus. Lines tangent to the curvature of the eye (red and blue) have been drawn for the flattest region of the eye and are used to calculate the curvature of the eye in the relevant axis (see methods and Bergman and Rutowski 2016 for more details). B) an anterior view of the flattest region of the eye used to calculate average facet diameter. *b / α* = eye surface length (*b)* in a given angle (*α), D* = facet diameter.

Using the lateral (for y curvature) or dorsal views (for x curvature), the localized curvature of the eye (*b* / *α*) was calculated in two axes via ImageJ (v. 1.50i; Schneider et al. 2012). First, we drew two lines perpendicular to the surface of the eye (Figure 1a; Bergman and Rutowski 2016). From these lines, *b* was calculated as the distance between the two points created by the intersection of the perpendicular lines to the eye edge, while the angular distance covered between these points (*α)* was calculated using the ImageJ angle function. These measurements were then combined (*b* / *α)* to calculate the distance of the eye’s surface covered in a given angle (*µm* per °).

Facet diameter (*D*) was calculated from two rows of ten facets in the flattest region of the eye (Figure 1B). Similar to studies in other Oedipodinae (Horridge, 1978), preliminary results found that facet diameter was relatively constant across the majority of the eye surface of each individual grasshopper.

Using the previous measurements, the inter-ommatidial angle (ΔΦ) in each axis was then calculated as:

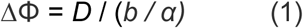

Where *D* = the facet diameter and *b / α* = eye surface length in a given angle.

Finally, visual acuity (VA, in degrees, smaller values indicate finer vision) in each axis was calculated as two times the inter-ommatidial angle:

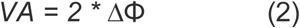

#### 2.1.3 Statistical Analysis

All data was analyzed in R version 3.4.4 (R CORE Team, 2018). To quantify the magnitude of the sexual size dimorphism, we first used a two-way ANOVA examining how total length varies with species and biological sex.

VA data was analyzed via two different approaches that produced similar results. First, we analyzed how VA in each axis was predicted by a variety of factors using generalized linear models (GLMs). We used the Akaike information criterion (AIC) to select the most parsimonious model(s). We treated models within 2 AIC as equally parsimonious. Second, because our GLM results suggested that biological sex and species were the main predictors of VA, we used two-way ANOVAs to examine how biological sex and species predict VA and its morphological determinants (eye curvature and facet size). Normality for all data was first checked using the Shapiro-Wilk normality test. In two cases data was either natural log transformed (VA_x_) or had an outlier excluded (vertical eye curvature) to restore normality. Student’s T-tests or Tukey HSDs were used for post-hoc analysis.

### 2.2 Experiment II: Eye size and facet count variation in Dissosteira carolina

#### 2.2.1 Study Organisms

To further explore the relationship between body size, eye size, and VA, we examined eye and ommatidia scaling within the species that showed the greatest body size dimorphism (*D. carolina*). Unfortunately logistical issues prevented us from returning to the same study population, so 48 *D. carolina* (n = 25 male and 23 female) were collected during the summer (June-September) of 2018 from a suburban site near manicured grass in Wooster, OH. Directly prior to imaging, grasshoppers were euthanized in a freezer for ∼1 hour. Weight (g), body length (head to tip of the thorax; *mm*), head size (*mm*), and biological sex were recorded for each individual.

#### 2.2.2 Eye size measurements in Dissosteira carolina

All individuals (25 males, 23 females) were photographed at a lower resolution (30x) then the previous experiment to ensure that all eyes completely fit within the images. Eye images were taken using an AmScope stereo trinocular microscope (United Scope LLC; Irvine, CA, USA) paired with a MU 1403 digital camera (United Scope LLC; MU1403; Irvine, CA, USA). Because band-winged grasshoppers have non-spherically shaped eyes, calibrated images of each eye were taken from three different angles (anterior, dorsal, lateral) and used to calculate eye size in all three axes (x, y, z) in ImageJ (v 1.8.0, Schneider et al. 2012). Note that this methodology only estimates the exposed eye depth in the z axis.

#### 2.2.3 Facet counts in Dissosteira carolina

To examine if sexes vary in number of receptors, we calculated the total number of facets per eye via eye castings of a subsample of the Ohio *D. carolina* population (e.g. Narendra et al., 2013). Individuals representative of the various eye sizes within each biological sex were used (n=3 of each sex). A single thin layer of #800 crystal clear nail polish (Sally Hansen Inc., New York, NY, USA) was applied to the entirety of the left eye and dried for 30-50 minutes. When dry, the eye castings were removed and cut horizontally and vertically into multiple sections with a #11 surgical scalpel blade (Swann-morton LTD., Sheffield, UK). Each section was then flattened between two microscope slides overnight before imaging at 30X magnification (Supplemental Figure 1) using an AmScope stereo trinocular microscope (United Scope LLC; Irvine, CA, USA) paired with a MU1403 digital camera (United Scope LLC; Irvine, CA, USA). Each eye section was assigned a non-identifying name, and facets were then counted by individuals blinded to original sex and grasshopper ID of the images. Sections originating from the same eye were then summed to determine the total facet count for each eye.

#### 2.2.4 Statistical Analysis

We examined how biological sex and eye axis vary via a two-way ANOVA in R (R CORE Team, 2018). Student’s T-tests or Tukey HSDs were used for post-hoc analysis.

## 3. RESULTS

### 3.1 Experiment I: VA variation by biological sex across three species

#### 3.1.1 Dimorphism in body size

Biological sex, species, and their interaction all significantly predicted total length in band-winged grasshoppers (Figure 2). Biological sexes significantly differed in total body length (p<0.001, df=1, F=257.93, two-way ANOVA), with females being on average 6.2-7.0 mm longer than their male counterparts (p<0.001, TukeyHSD). Additionally, all three species significantly differed from each other in total length (p<0.001, df=2, F=482.81, two-way ANOVA; all TukeyHSD p<0.001). The interaction between biological sex and species on total body length was also significant (p<0.001, df=2, F=12.71), with *Dissosteira carolina* showing the largest dimorphism.

**Figure 2:**
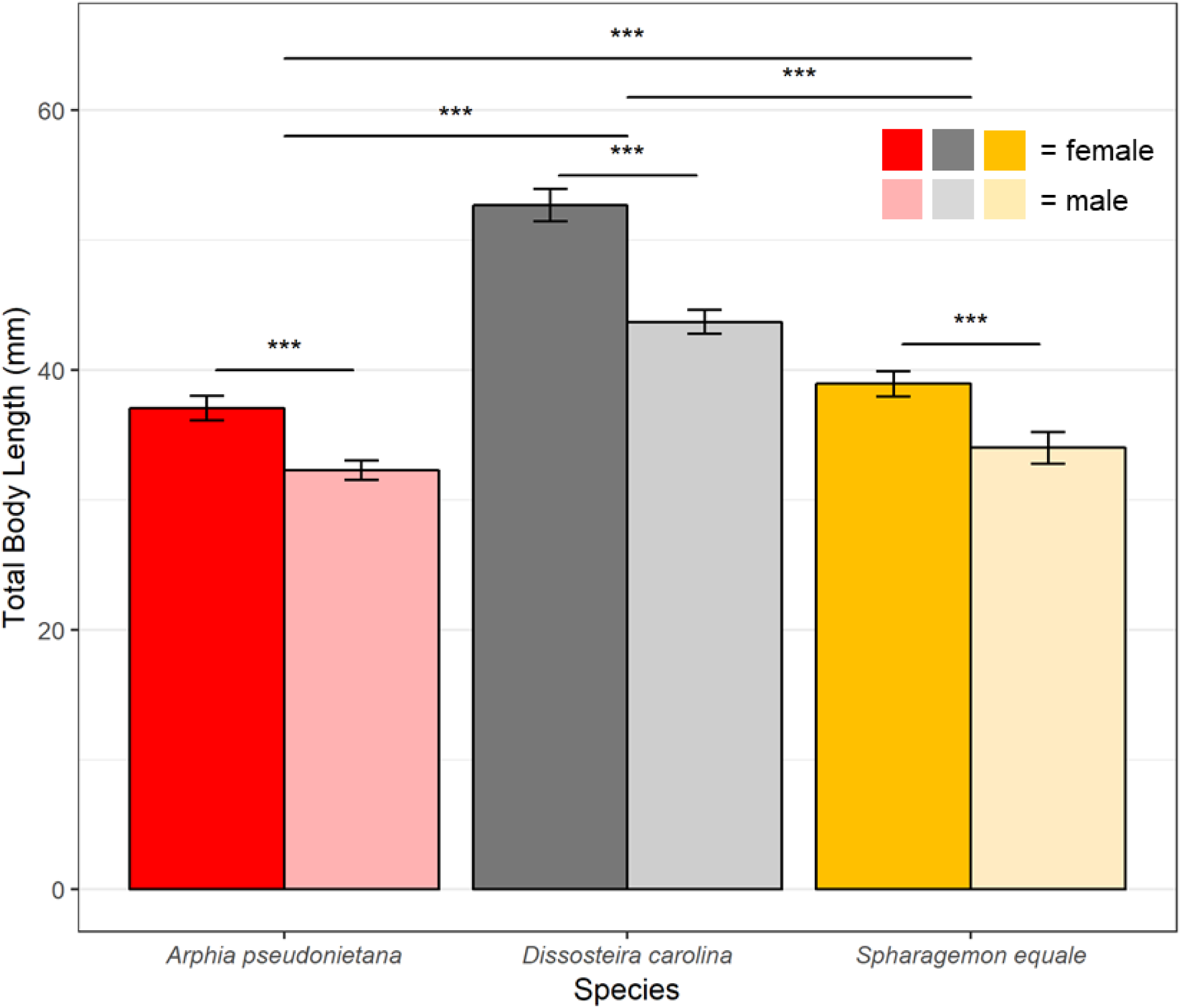
Body length in three species of band-winged grasshoppers. All three species show a sexual size dimorphism, with biological females being longer than males (p<0.001, df=1, F=257.93, two-way ANOVA). Additionally, total length varies between species, with all three showing significant differences from each other (p<0.001 in all comparisons, TukeyHSD). There is also a significant interaction between biological sex and species on total body length (p<0.001, df=2, F=12.71). Sample sizes (from left to right) are 18, 18, 16, 16, 15, and 15 individuals respectively. Error bars indicate 95% CI, significance symbols are based on post -hoc analyses (Student’s T-test for biological sex differences within a species, TukeyHSD for species differences).

#### 3.1.2 Visual acuity perpendicular to the horizon (VA_y_)

Visual acuity varied depending on the axis of view in all three species; amongst all individuals measured (n=98) visual acuity perpendicular to the horizon (VA_y_; average of 2.17°) was twice as sharp as visual acuity parallel to the horizon (VA_x_; average of 4.34°; Figure 3). As such, results are presented separately for each axis of view.

**Figure 3:**
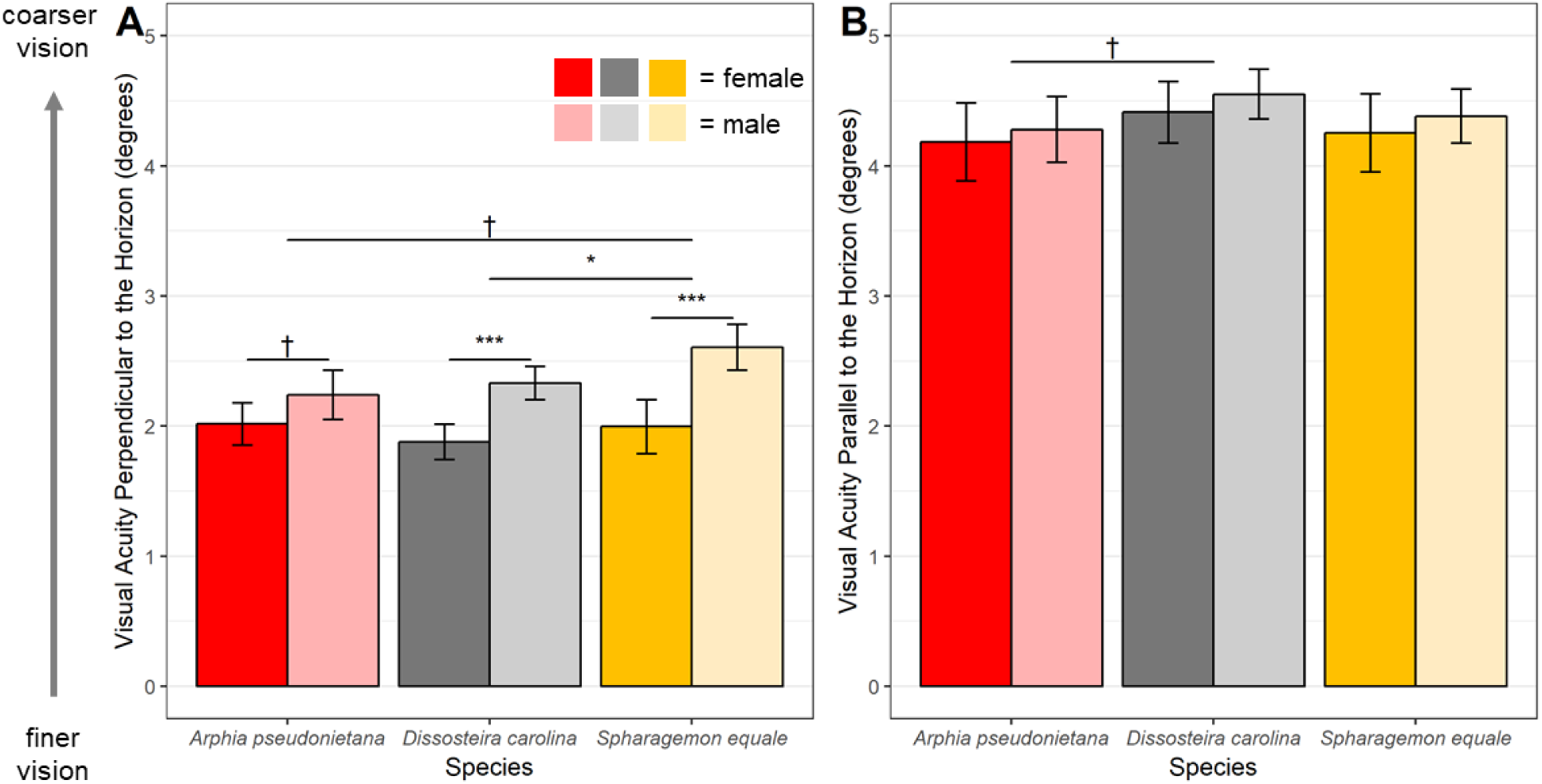
Visual acuity in the two axes of vision in band-winged grasshoppers. Visual acuity in band-winged grasshoppers is sexually dimorphic, but only in one axis. Units are presented in degrees, so finer vision is indicated by smaller values. **A)** Biological females typically have finer visual acuity than males in the axis perpendicular to the horizon (p<0.001, df=1, F=41.04, two-way ANOVA). Additionally, species differ significantly from one another (p<0.05, df=2, F=3.44) with *D. carolina* having finer vision than *S. equa*le (p<0.05, TukeyHSD). **B)** In contrast to A, visual acuity parallel to the horizon is both coarser and shows no significant sexual dimorphism or species-specific effect (see text for details). **A-B)** Sample sizes (from left to right) are 18, 18, 16, 16, 15, and 15 individuals respectively in each panel. Error bars indicate 95% CI, significance symbols are based on post-hoc analyses (Student’sT-test for biological sex differences within a species, TukeyHSD for species differences).

The most parsimonious models of VA_y_ all included biological sex, species, and their interactions as factors (Table 1). Culmulatively, models including both sex and species as predictors accounted for 92% of relative model probability (*w*_*i*_). Some equally parsimonious models included size measurments as well, but these did not significanty improve the model (Table 1). In the GLM including biological sex and species, males were estimated to have ∼0.416 degrees coarser vision (Std error=0.066, t=6.275, p<0.0001, GLM).

**Table 1.** Summary of predictor combinations explaining visual acuity perpendicular to the horizon (VA_y_)

| Model | $\Delta AIC$ | $I_i$ | $w_i$ |
| --- | --- | --- | --- |
| **$VA_y \sim \text{Sex} * \text{Species} + \text{Length}$** | **0** | **1.00** | **0.28** |
| **$VA_y \sim \text{Sex} * \text{Species}$** | **0.1** | **0.95** | **0.27** |
| **$VA_y \sim \text{Sex} * \text{Species} + \text{Weight}$** | **1.3** | **0.52** | **0.15** |
| $VA_y \sim \text{Sex} + \text{Species} + \text{Length}$ | 2.2 | 0.33 | 0.09 |
| $VA_y \sim \text{Sex} + \text{Species}$ | 2.3 | 0.32 | 0.09 |
| $VA_y \sim \text{Sex} + \text{Species} + \text{Weight}$ | 3.7 | 0.16 | 0.04 |
| $VA_y \sim \text{Sex} + \text{Length}$ | 4.5 | 0.11 | 0.03 |
| $VA_y \sim \text{Sex} + \text{Weight}$ | 4.6 | 0.10 | 0.03 |
| $VA_y \sim \text{Sex}$ | 5 | 0.08 | 0.02 |
| $VA_y \sim \text{Weight}$ | 15.3 | 0.00 | 0.00 |
| $VA_y \sim \text{Length}$ | 24.3 | 0.00 | 0.00 |
| $VA_y \sim \text{Species}$ | 34.6 | 0.00 | 0.00 |

$\Delta AIC$ values were calculated relative to the best fit model. Models in bold are equally parsimonious (within 2 $\Delta AIC$ ). $I_i$ = model's relative likelihood, $w_i$ = model probability

Results from a two-way ANOVA mirrored those of the GLM (Figure 3A). VA_y_ was dependent on both biological sex (p<0.001, df=1, F=41.04, two-way ANOVA) and species (p<0.05, df=2, F=3.44, two-way ANOVA). Across all individuals measured, female VA_y_ (average = 1.96°) was estimated to be 21% finer (p<0.001, 95% CI = 0.29°-0.55°, TukeyHSD) than male VA_y_ (average = 2.38°; Figure 2A). Additionally, the large *D. carolina* had significantly finer VA_y_ than *S. equale* (p<0.05, 95% CI= 0.002°-0.39°, TukeyHSD), however other species comparisons did not vary significantly (all p<0.05, TukeyHSD, Figure 2A). The interaction between sex and species on VA_y_ trended towards significance (p=0.055, df=2, F=2.994, two-way ANOVA). Within any biological sex and species combination examined, size did not correlate with VA_y_ (all r^2^ < 0.2, all p>0.1; Supplemental Figure 2), suggesting that differences in size may act between the sexes but not within.

#### 3.1.3 Visual acuity parallel to the horizon (VA_x_)

In contrast to VA_y_, VA_x_ showed no sexual dimorphism nor species-specific differences. VA_x_ was not normally distributed (p<0.01, W = 0.95755, Shapiro-Wilk normality test), and therefore a natural log transformation was used to restore normality. Neither biological sex (p=0.18, df=1, F=1.86, two-way ANOVA), species (p=0.07, df=2, F=2.73, two-way ANOVA), nor their interactions (p=0.99, df=2, F=0.013, two-way ANOVA) were significant predictors of the natural log of VA_x_ (Figure 3B).

#### 3.1.4 Dimorphism in the Morphological Determinants of VA

At the morphological level, compound eyes can achieve finer VA if they have either 1) a flatter curvature or 2) smaller facets. In female band-winged grasshoppers, changes in eye curvature --- and not facet size --- led to finer VA_y_ (Figure 4). Vertical eye curvature was not normally distributed (p<0.01, W=0.95, Shapiro-Wilk normality test) because of the presence of one female *S. equale* outlier with a particularly flat eye (perpendicular eye curvature = 51 µm/degree). Removal of the outlier restored normality (p=0.18, W=0.98, Shapiro-Wilk normality test). Even with this especially flat eye removed, females still have a vertical curvature that is ∼18% flatter than their male counterparts (p<0.001, df=1, F=49.56, two-way ANOVA, Figure 4A). Additionally, there are species-specific differences in vertical eye curvature (p<0.001, df=2, F=14.58, two-way ANOVA) with post-hoc testing revealing that the larger *D. carolina* has a flatter vertical eye curvature than either of the two other species (p<0.001 in both cases, TukeyHSD). The interactions between biological sex and species on vertical eye curvature were trending towards significance (p=0.06, df=2, F=2.87), suggesting that the dimorphism in vertical curvature may vary between species.

**Figure 4:**
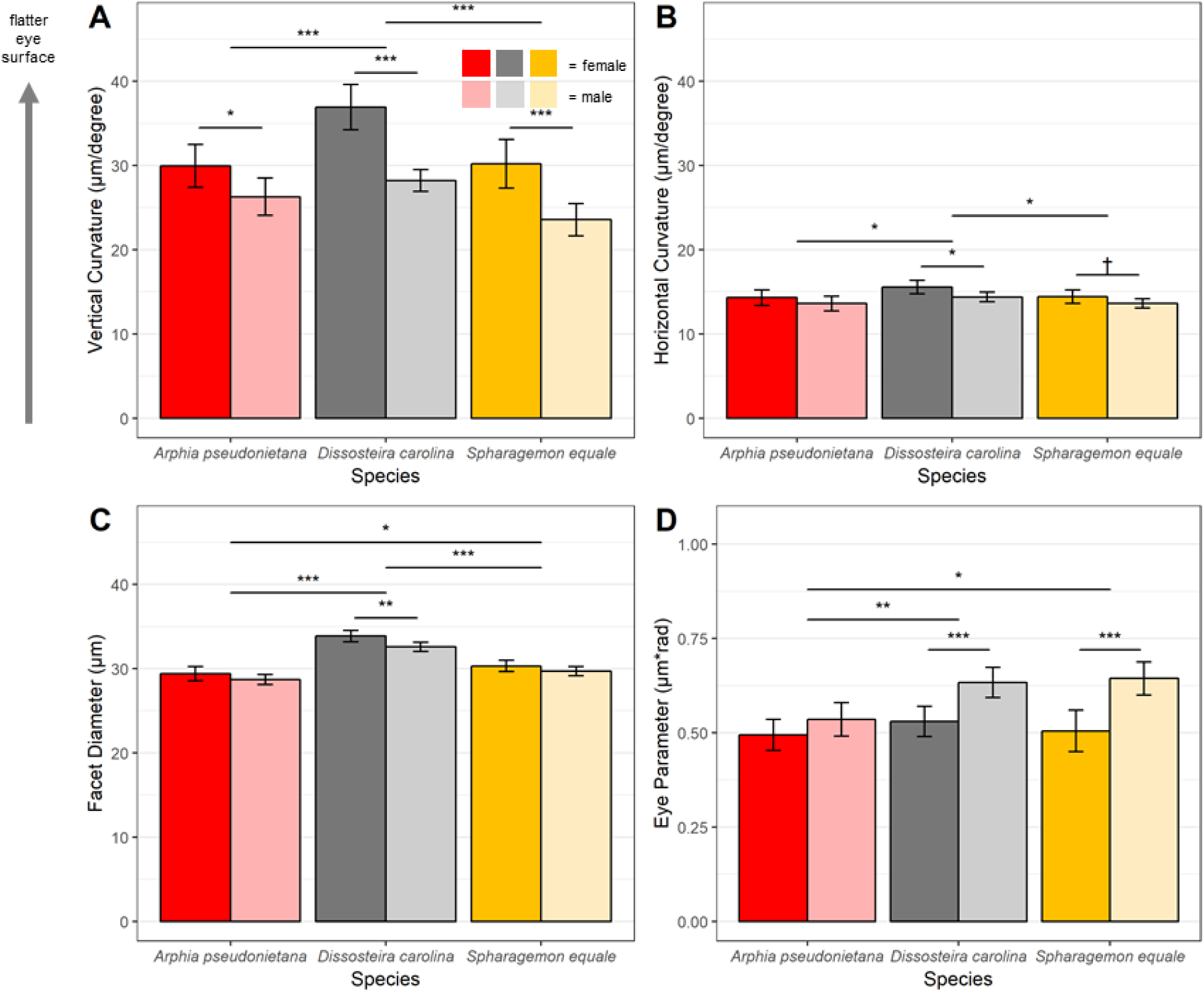
Morphological determinants of visual acuity in band-winged grasshoppers. Eye curvature is the biggest morphological driver of the sexually dimorphic visual acuity in these three species of band-winged grasshoppers. **A)** Biological females had ∼18% flatter vertical eye curvature in the region of the eye with the sharpest visual acuity (p<0.001, df=1, F=49.6, two-way ANOVA). **B)** Although significant, the differences in horizontal curvature were not as pronounced, as female curvature was only ∼6.5% flatter than males (p<0.01, df=1, F=8.65, two-way ANOVA). **C)** Female facet diameters were slightly larger (∼2.8%) than male facet diameters (p<0.01, df=1, F=1.52, two-way ANOVA). **D)** The observed eye parameter values are typical of diurnal insects and varied significantly between both sexes (p<0.001, df=1, F=29.15, two-way ANOVA) and species (p<0.001, df=2, F=2.92, two-way ANOVA). **A-D)** Sample sizes (from left to right) in each panel are 17, 18, 16, 16, 15, and 15 individuals respectively. Error bars indicate 95% CI, significance symbols are based on post-hoc analyses (Student’sT-test for biological sex differences, TukeyHSD for species differences).

Similarly to VA_x_ and VA_y_, differences in eye curvature between the sexes were less pronounced in the horizontal axis than in the vertical axis (Figure 4B). Both biological sex (p<0.01, df=1, F=8.65, two-way ANOVA) and species (p<0.01, df=2, F=4.85) significantly affected horizontal eye curvature. However, female horizontal curvature was only ∼6.5% flatter than males. There was no significant effect of the interaction between biological sex and species on horizontal eye curvature (p=0.78, df=2, F=0.252).

Females had slightly larger facet diameters than males (∼0.9 µm larger or 2.8%, Figure 4C) across all three species measured (p<0.01, df=1, F=11.47, two-way ANOVA). Additionally, all three species differed significantly in facet size (p<0.001, df=2, F=96.06, two-way ANOVA) with the larger *D. carolina* having the largest facets (all p<0.05, Tukey HSD). There was no significant effect of the interaction between biological sex and species on facet diameter (p=0.52, df=2, F=0.651, two-way ANOVA). The combination of changes in receptor spacing and facet size led to an eye parameter (Figure 4D) that varied significantly between both sex (p<0.001, df=1, F=29.15, two-way ANOVA) and species (p<0.001, df=2, F=2.92, two-way ANOVA). The interactions between biological sex and species on the eye parameter were trending towards significance (p=0.059, df=2, F=2.918).

### 3.2 Experiment II: Eye size and facet count variation in Dissosteira carolina

Further investigation of the VA_y_ dimorphism in the species that showed the greatest sexual size dimorphism (*D. carolina*) found that females typically have larger eyes than males, but not in every axis (Figure 5). Compared to males, females had significantly larger eye diameters in the horizontal (X; ∼13% increase) and vertical (Y, ∼23% increase) axes, but did not vary in exposed eye depth (Z; p<0.001, df=1, F=28.90, two-way ANOVA; sex*interaction p<0.001, df=2,F=7.98). Additionally, both male and female eyes varied significantly in diameter between the axes (p<0.001, df=2, F=240.73).

**Figure 5:**
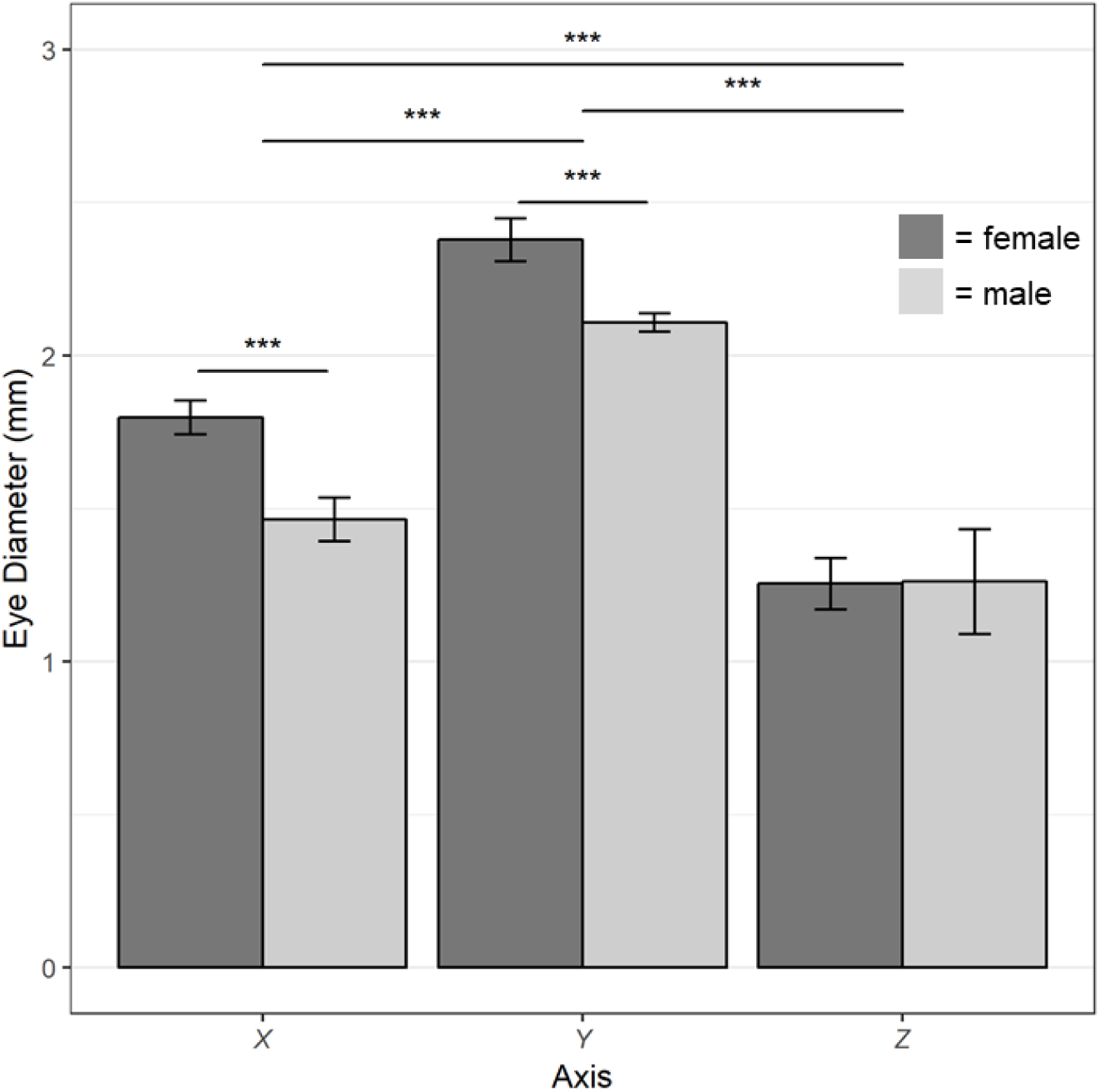
Eye size in the band-winged grasshopper *Dissosteira carolina*. Eye size is both asymmetrical and varies between the sexes. Females have larger eyes in the X and Y axis, but not in exposed eye depth (Z; p<0.001, df=1, F=28.90, two-way ANOVA). Exposed eye depth (Z) only measures exposed eye surface and may be an underestimate of total size. Sample size are 23 females and 25 males. Error bars indicate 95% CI, significance symbols are based on post-hoc analyses (Student’s T-test for biological sex differences, TukeyHSD for species differences).

As expected based on differences of eye size and facet diameter, facet counts of a subset of representative individuals suggest that the female eyes have more facets than males (Table 2). Although there was variability in the number of facets seen in each sex, representative females (average = 5565 facets, n=3) had eyes with ∼19% more facets than males (4679 facets, n=3).

**Table 2:** Whole eye facet counts in representative *Dissosteira carolina* individuals

| eye size percentile | female | male |
| --- | --- | --- |
| ~25th | 4901 facets | 4571 facets |
| ~50th | 5514 facets | 4453 facets |
| ~75th | 6280 facets | 5013 facets |

## 4. Discussion

Most band-winged grasshoppers have non-spherical eyes featuring an elongated vertical axis that gives them a jellybean-like shape. Similar to other invertebrates with non-spherical eyes (Bagheri et al., 2020; Kelber and Somanathan, 2019) this leads to better visual acuity in the axis of elongation: in band-winged grasshoppers visual acuity in the vertical axis (VA_y_) is around twice as fine as that in the horizontal axis (VA_x_; Figure 3). VA_y_ appears to be enhanced by a particularly flat vertical region near the center of the eye.

VA_y_ in this particularly flat region of the eye is sexually dimorphic, resulting in female grasshoppers --- the larger of the two sexes --- having finer vision (Figure 3a). Similar to other grasshopper species (Hochkirch and Gröning, 2008; Otte, 1984), female band-winged grasshoppers were substantially larger than males in this study (Figure 2). Similarly VA_y_ --- but not VA_x_ --- was sexually dimorphic, with females having ∼21% finer vision than males in the sharpest region of the eye (Figure 3a-b). This VA_y_ dimorphism is similar in magnitude to the classic “love spot” seen in male flies (although without the accompanying sensitivity specializations, (Land and Eckert, 1985)) however it is in the opposite direction: females have finer vision rather than males. This suggests that much like we observe between species (Caves et al., 2017; Caves et al., 2018; Kiltie, 2000; Land and Nilsson, 2012; Veilleux and Kirk, 2014), size differences can lead to finer vision within a species. Notably, within either sex there was no further effect of size on VA_y_ (Supplemental Figure 2). Thus, in some cases the positive relationship between acuity and size could be constrained to individuals with consistently different developmental processes such as sex and/or morph differences.

The magnitude of the VA_y_ dimorphism varied by species as all of the top predictor models included a term for the interaction between sex and species (Table 1). This was driven by *Arphia pseudonietana* showing a much smaller dimorphism than either of the two other species (Figure 6) that when considered by itself was only trending towards significantly varying between the sexes (Figure 3A). Notably, *A. pseudonietana* was one of the species that showed a smaller size dimorphism (Figure 2), which could lead to a less pronounced VA dimorphism. However, *Spharagemon equale* has a similar size dimorphism but showed a substantial VA_y_ dimorphism suggesting that selective pressures and/or developmental constraints on band-winged vision could be acting differently across species.

**Figure 6:**
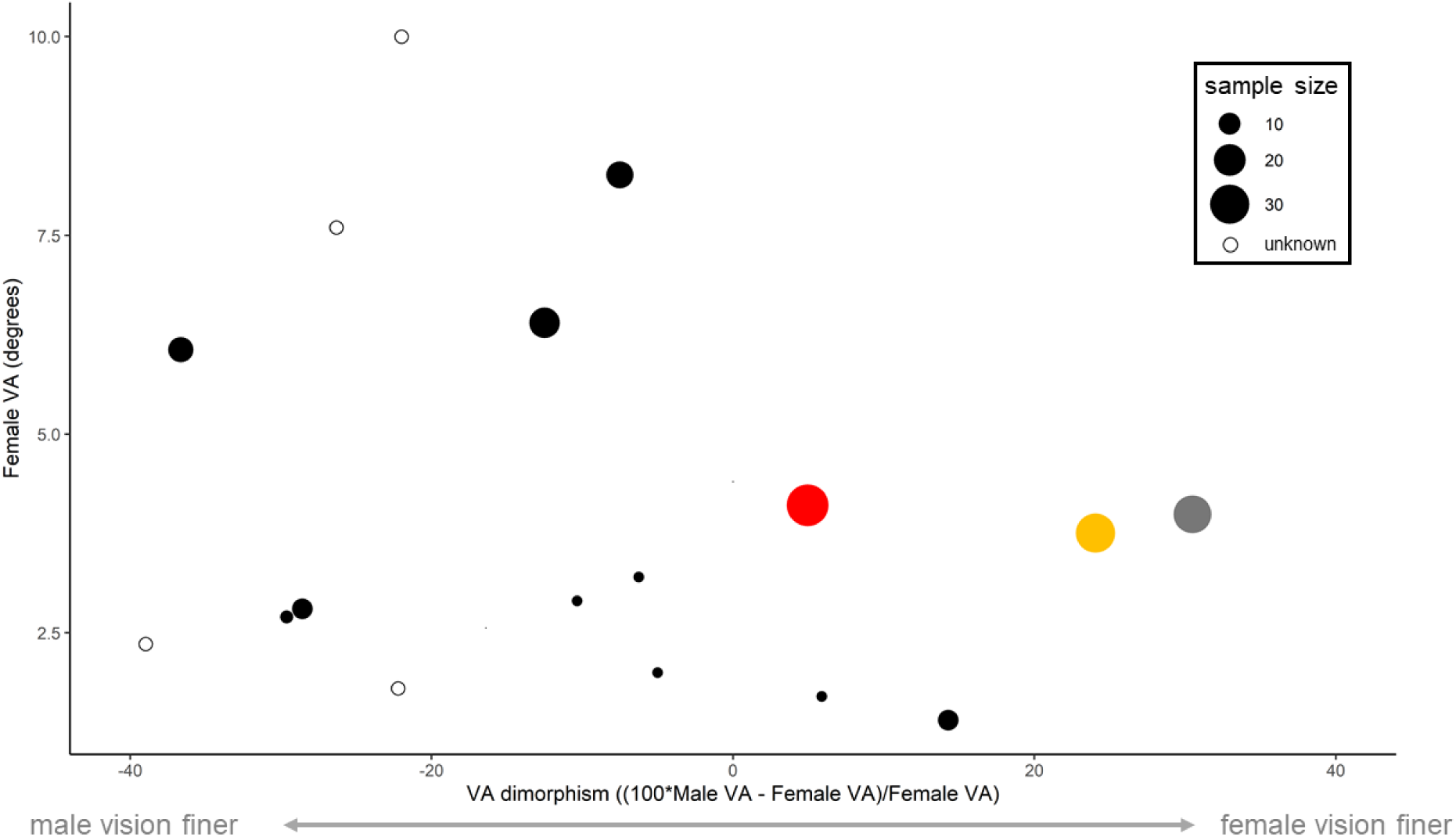
In insects with moderate spatial vision (VA < 10°), studies that report VA separately for each sex have typically 1) not shown females with finer vision than males and 2) been based on smaller sample sizes. Our study (colored circles) is one of the first to suggest a female-biased VA dimorphism in insects with the potential for Type I spatial vision (VA < 10°). Red = *Arphia pseudonietana*, Yellow = *Spharagemon equale*, gray = *Dissosteira carolina*. Data and references can be found in Supplemental Table 1.

Across all three species examined, the dimorphism was morphologically caused by a flatter eye surface and not a reduction in facet diameter (Figure 4). Although both changes can lead to finer acuity, decreasing facet diameter is not typically seen in insects because it may also reduce overall sensitivity (Kirschfeld, 1976; Land, 1997). The acute zones of many insects require both fine VA and heightened sensitivity (Warrant, 2016), and therefore often feature flat regions with large facets. In band-winged grasshoppers, females either had similar or slightly larger facets than males. However, their eye’s flatter curvature led to a substantial VA dimorphism. As a result, female band-winged grasshoppers have finer VA_y_ without sacrificing sensitivity by decreasing facet diameter (Figure 4).

Further study in *D. carolina* suggests that the sexual size dimorphism corresponds with both an eye size dimorphism (Figure 5) and a facet number dimorphism (Table 1). Altogether, this suggests that female *D. carolina* improve VA_y_ by having larger eyes with more facets. Surprisingly, we also found that females had a larger eye diameter in the horizontal axis, despite showing no improvement in VA_x_ compared to males. This increase in X eye diameter suggests that female *D. Carolina* may improve VA, sensitivity, and/or field of view in other regions of their eye that were not measured within this study. Future studies should further explore the regional variation within female and male eyes to elucidate how visual performance changes outside of the central region of the eye.

It is unknown what --- if any --- selective advantage female band-winged grasshoppers may gain from having larger eyes with finer vision. One possibility is that females could benefit from finer vision by more accurately interpreting visual signals. Although mating systems and behaviors are variable across the subfamily, many species use a variety of potential visual signals including those involving either their colorful hindwings or leg movements (Kerr, 1974; Otte, 1970; Otte, 1984; Willey and Willey, 1969). Females may therefore benefit from finer vision by being better able to perceive and interpret these potential signals of species identity and/or mate quality. Additionally, females could use their enhanced vision to initiate anti-predator defenses at a greater distance. Band-winged grasshoppers have a suite of defensive behaviors in response to approaching predators (Collier and Hodgson, 2017; Santer et al., 2012). Initiating these defenses at greater distances could be especially beneficial to females because they may take a longer time than males to reach sexual maturity even once in their final adult instar (Pfadt, 1994). Alternatively, the VA dimorphism could be a byproduct of increased body size and serve no beneficial function. Future studies utilizing the natural variation in band-winged size dimorphism, mating systems, and development could elucidate how these factors contribute to the visual dimorphism seen in this study.

Surprisingly, our results are one of the first studies to show females with finer vision than males in species with the potential for high-resolution vision (Figure 6). In insects, females of the tiny parasitic wasp *Trichogramma evanescens* have larger eyes with ∼23% finer vision than males. However, because of their small size, female VA (∼20°) is much coarser than what was measured in this study and likely limits their visually guided behaviors (Nilsson, 2009; Nilsson, 2013). In mammals, Seymoure and Juraska (1997) found that female rats behaviorally outperform males for coarse stimuli but that this sex difference disappeared at finer stimuli. And in fish, Corral-López et al. (2017) found that female artificially selected guppies have finer VA than males. However, unlike what we report in this study, the difference disappears when controlling for body size.

We believe that the documentation of females with finer vision than males is lacking— not because it is a rare phenomenon—but rather little research has so far been conducted on the topic. Aside from the well documented love-spot in species of flying insects, few visual acuity studies have examined differences between the sexes and studies often do not report the sexes of the individuals measured. Instead most studies have prioritized either an ecological approach [sampling only a few individuals because they sample many species; *e.g*. (Collin and Partridge, 1996; Lisney et al., 2012; Pettigrew et al., 1988)] or a retinal topography approach [sampling only a few individuals because of the extensive work it takes in each individual; *e.g*. (Coimbra et al., 2013; Landgren et al., 2014) making extensive comparisons between the sexes not possible. More recent studies in insects have found variation in eye-scaling within insect species but have so far focused on the morphs of eusocial insects (Perl and Niven, 2016a; Perl and Niven, 2016b; Taylor et al., 2019). Because eye size is a major factor influencing visual performance (Cronin et al., 2014; Kirschfeld, 1976; Land and Nilsson, 2012), body size increases that lead to eye size increases could be utilized for VA improvements (Corral-López et al., 2017). Based on the prevalence of sexual size dimorphisms throughout many animal groups (Kuntner and Coddington, 2020; Lislevand et al., 2007; Parker, 1992; Teder, 2014), VA differences between the sexes may be an under-documented --- yet not uncommon --- phenomenon that warrants further consideration and exploration.

## 5. Acknowledgements

The authors thank Dr. Carrie Veilleux and Dr. Eleanor Caves for comments on an earlier draft of this manuscript. The authors would also like to thank Cesar Nufio for initial help with species identification. Eran Maina, Jack Redick, Jack Whalen, and Miura Wiley all provided help in the field during experiment II.

## Literature Cited

Bagheri, Z. M., Jessop, A.-L., Kato, S., Partridge, J. C., Shaw, J., Ogawa, Y. and Hemmi, J. M. (2020). A new method for mapping spatial resolution in compound eyes suggests two visual streaks in fiddler crabs. The Journal of Experimental Biology 223, jeb210195.

Barlow, H. B. (1952). The size of ommatidia in apposition eyes. Journal of Experimental Biology 29, 667–674.

Bergman, M. and Rutowski, R. L. (2016). Eye morphology and visual acuity in the pipevine swallowtail (*Battus philenor*) studied with a new method of measuring interommatidial angles. Biological Journal of the Linnean Society 117, 646–654.

Caves, E. M., Sutton, T. T. and Johnsen, S. (2017). Visual acuity in ray-finned fishes correlates with eye size and habitat. J. Exp. Biol. 220, 1586.

Caves, E. M., Brandley, N. C. and Johnsen, S. (2018). Visual acuity and the evolution of signals. Trends in Ecology & Evolution 33, 358–372.

Caves, E. M., Nowicki, S. and Johnsen, S. (2019). Von uexküll revisited: addressing human biases in the study of animal perception. Integrative and Comparative Biology 59, 1451–1462.

Coimbra, J. P., Hart, N. S., Collin, S. P. and Manger, P. R. (2013). Scene from above: Retinal ganglion cell topography and spatial resolving power in the giraffe (*Giraffa camelopardalis*). Journal of Comparative Neurology 521, 2042–2057.

Collier, A. and Hodgson, J. Y. S. (2017). A shift in escape strategy by grasshopper prey in response to repeated pursuit. Southeastern Naturalist 16, 503–515.

Collin, S. P. and Partridge, J. C. (1996). Retinal specializations in the eyes of deep-sea teleosts. Journal of Fish Biology 49, 157–174.

Corral-López, A., Garate-Olaizola, M., Buechel, S. D., Kolm, N. and Kotrschal, A. (2017). On the role of body size, brain size, and eye size in visual acuity. Behavioral Ecology and Sociobiology 71, 179.

Cronin, T. W., Johnsen, S., Marshall, N. J. and Warrant, E. J. (2014). Visual Ecology. STU-Student edition. Princeton University Press.

Hochkirch, A. and Gröning, J. (2008). Sexual size dimorphism in Orthoptera (*sens. str*.) — a review. Journal of Orthoptera Research 17, 189–196.

Horridge, G. A. (1978). The separation of visual axes in apposition compound eyes. Philosophical Transactions of the Royal Society of London. B, Biological Sciences 285, 1–59.

Jander, U. and Jander, R. (2002). Allometry and resolution of bee eyes (Apoidea). Arthropod Structure & Development 30, 179–193.

Jordan, L. A. and Ryan, M. J. (2015). The sensory ecology of adaptive landscapes. Biol Lett 11,.

Kelber, A. and Somanathan, H. (2019). Spatial vision and visually guided behavior in apidae. Insects 10, 418.

Kerr, G. E. (1974). Visual and acoustical communicative behaviour in *dissosteira Carolina* (orthoptera: acrididae). The Canadian Entomologist 106, 263–272.

Kiltie, R. A. (2000). Scaling of visual acuity with body size in mammals and birds. Functional Ecology 14, 226–234.

Kirschfeld, K. (1976). The Resolution of Lens and Compound Eyes. In Neural Principles in Vision (ed. Zettler, F.) and Weiler, R.), pp. 354–370. Berlin, Heidelberg: Springer.

Kirschfeld, K. and Wenk, P. (1976). The Dorsal Compound Eye of Simuliid Flies: Zeitschrift für Naturforschung C 31, 764–765.

Kuntner, M. and Coddington, J. A. (2020). Sexual Size Dimorphism: Evolution and Perils of Extreme Phenotypes in Spiders. Annual Review of Entomology 65, 57–80.

Land, M. F. (1997). Visual Acuity in Insects. Annual Review of Entomology 42, 147–177.

Land, M. F. and Eckert, H. (1985). Maps of the acute zones of fly eyes. J. Comp. Physiol. 156, 525–538.

Land, M. F. and Nilsson, D.-E. (2012). Animal Eyes. Oxford University Press.

Landgren, E., Fritsches, K., Brill, R. and Warrant, E. (2014). The visual ecology of a deep-sea fish, the escolar *Lepidocybium flavobrunneum* (Smith, 1843) <sup/>. Philosophical Transactions of the Royal Society B: Biological Sciences 369, 20130039.

Lislevand, T., Figuerola, J. and Székely, T. (2007). Avian body sizes in relation to fecundity, mating system, display behavior, and resource sharing: *ecological archives* e088-096. Ecology 88, 1605–1605.

Lisney, T. J., Iwaniuk, A. N., Kolominsky, J., Bandet, M. V., Corfield, J. R. and Wylie, D. R. (2012). Interspecifc variation in eye shape and retinal topography in seven species of galliform bird (Aves: Galliformes: Phasianidae). Journal of Comparative Physiology A 198, 717–731.

Narendra, A., Alkaladi, A., Raderschall, C. A., Robson, S. K. A. and Ribi, W. A. (2013). Compound eye adaptations for diurnal and nocturnal lifestyle in the intertidal ant, polyrhachis sokolova. PLOS ONE 8, e76015.

Nilsson, D.-E. (2009). The evolution of eyes and visually guided behaviour. Philosophical Transactions of the Royal Society B: Biological Sciences 364, 2833–2847.

Nilsson, D.-E. (2013). Eye evolution and its functional basis. Visual Neuroscience 30, 5–20.

Otte, D. (1970). A comparative study of communicative behavior in grasshoppers. Miscellaneous Publications: Museum of Zoology, University of Michigan.

Otte, D. (1984). The North American Grasshoppers, Volume 2, Acridiae, Oedipodinae. Cambridge (MA): Harvard University Press.

Parker, G. A. (1992). The evolution of sexual size dimorphism in fish*. Journal of Fish Biology 41, 1–20.

Partan, S. and Marler, P. (2002). The Umwelt and its relevance to animal communication: Introduction to special issue. Journal of Comparative Psychology 116, 116–119.

Perl, C. D. and Niven, J. E. (2016a). Differential scaling within an insect compound eye. Biology Letters 12, 20160042.

Perl, C. D. and Niven, J. E. (2016b). Colony-level differences in the scaling rules governing wood ant compound eye structure. Sci Rep 6, 1–8.

Pettigrew, J. D., Dreher, B., Hopkins, C. S., McCall, M. J. and Brown, M. (1988). Peak density and distribution of ganglion cells in the retinae of microchiropteran bats: implications for visual acuity. Brain, Behavior and Evolution 32, 39–56.

Pfadt, R. E. (1994). Field Guide to Common Western Grasshoppers. Wyoming Agricultural Experiment Station.

R CORE Team (2018). R: A Language and Environment for Statistical Computing. Vienna, Austria: R Foundation for Statistical Computing.

Rutowski, R. L., Gislén, L. and Warrant, E. J. (2009). Visual acuity and sensitivity increase allometrically with body size in butterflies. Arthropod Structure & Development 38, 91–100.

Santer, R. D., Rind, F. C. and Simmons, P. J. (2012). Predator versus prey: locust looming-detector neuron and behavioural responses to stimuli representing attacking bird predators. PLoS ONE 7, e50146.

Seymoure, P. and Juraska, J. M. (1997). Vernier and grating acuity in adult hooded rats: The influence of sex. Behavioral Neuroscience 111, 792–800.

Spaethe, J. and Chittka, L. (2003). Interindividual variation of eye optics and single object resolution in bumblebees. Journal of Experimental Biology 206, 3447–3453.

Taylor, G. J., Tichit, P., Schmidt, M. D., Bodey, A. J., Rau, C. and Baird, E. (2019). Bumblebee visual allometry results in locally improved resolution and globally improved sensitivity. eLife 8, e40613.

Teder, T. (2014). Sexual size dimorphism requires a corresponding sex difference in development time: a meta-analysis in insects. Functional Ecology 28, 479–486.

Uexküll, J. von (2013). A Foray into the Worlds of Animals and Humans: with A Theory of Meaning. U of Minnesota Press.

Veilleux, C. C. and Kirk, E. C. (2014). Visual acuity in mammals: effects of eye size and ecology. Brain, Behavior and Evolution 83, 43–53.

Warrant, E. J. (2016). Sensory matched filters. Current Biology 26, R976–R980.

Willey, R. B. and Willey, R. L. (1969). Visual and acoustical social displays by the grasshopper arphia conspersa (orthoptera: acrididae). Psyche: A Journal of Entomology 76, 280–305.

