## Supplemental Information for "A sexual dimorphism in the spatial vision of band-winged grasshoppers"

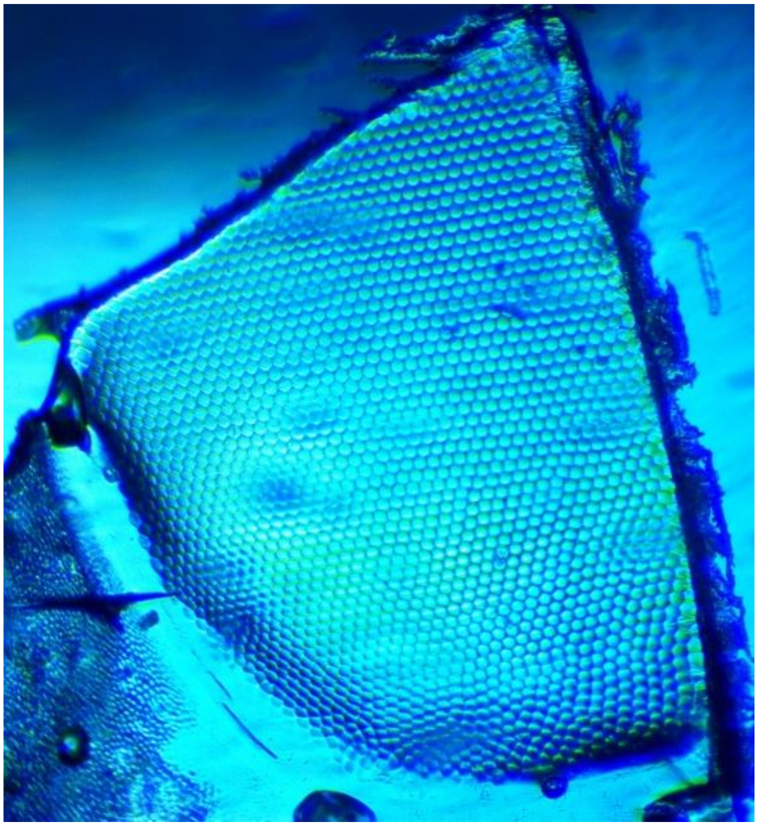

**Supplemental Figure 1:** Representative eye casting section used in calculating total number of facets in *Dissosteira carolina*. Number of facets were counted by individuals blinded to the original sex and grashopper ID. Sections from the same grasshopper were then summed to estimate the eye's total number of facets.

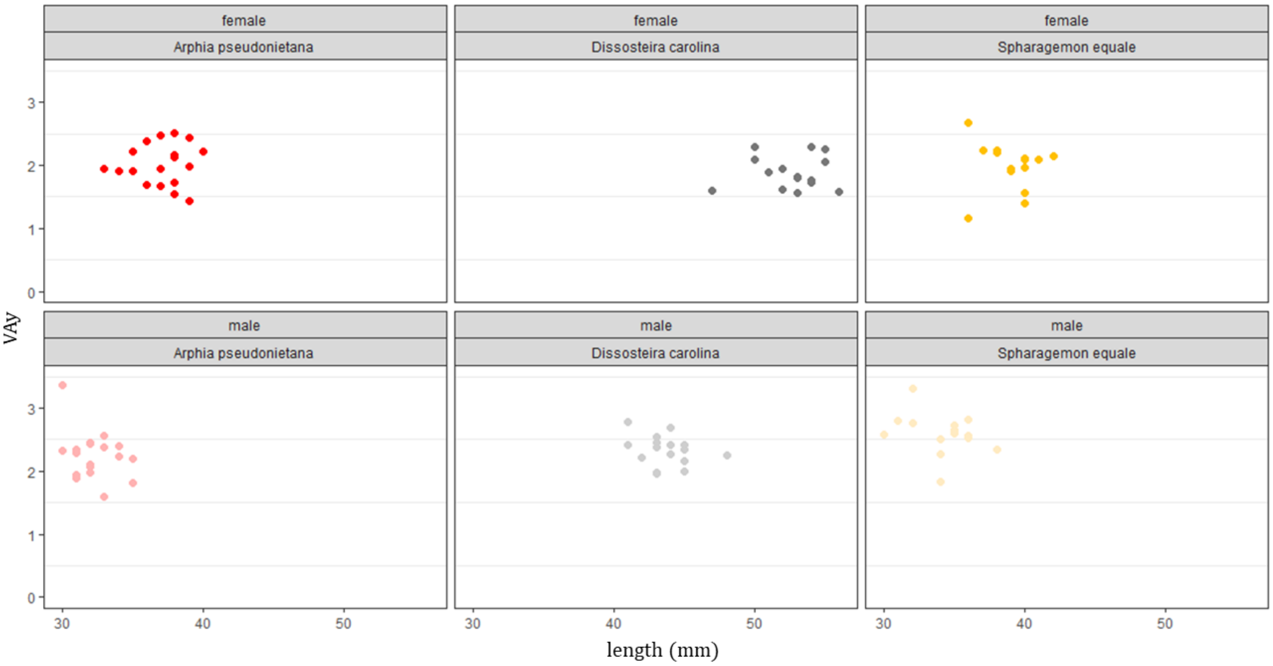


**Supplemental Figure 2: The relationship between size and $VA_y$ within all biological sex and species combinations.** Although there was variation within each sex and size class, size was not significantly correlated with $VA_y$ in any case examined. $r^2$, estimated slope ± standard error, and p-value for slope differs from zero are as follows: *Arphia pseudonietana* female ($r^2$ = 0.003, slope = 0.01 ± 0.04, p = 0.83), *Dissosteira carolina* female ($r^2$ = 0.001, slope = 0.003 ± 0.03, p = 0.92), *Spharagemon equale* female ($r^2$ = 0.01, slope = -0.02 ± 0.06, p = 0.72), *Arphia pseudonietana* female ($r^2$ = 0.11, slope = -0.08 ± 0.06, p = 0.18), *Dissosteira carolina* male ($r^2$ = 0.08, slope = 0.04 ± 0.03, p = 0.29), *Spharagemon equale* male ($r^2$ = 0.1, slope = -0.05 ± 0.04, p = 0.24).

| Supplemental Table 1: Data and References for Figure 6 | | | | | | | |
|---|---|---|---|---|---|---|---|
| Species | VA dimorphism (100 * (M - F)/F) | female VA | male VA | n (F) | n (M) | n (total) | citation |
| *Acentria ephemerella* | -7.5 | 8.3 | 7.6 | 8 | 8 | 16 | [1] |
| *Araschinia levana* | -6.3 | 3.2 | 3 | 2 | 2 | 4 | [2] |
| *Arphia pseudonietana* | 5 | 4.1 | 4.3 | 18 | 18 | 36 | this study |
| *Asterocampa leilia* | -28.6 | 2.8 | 2 | 5 | 5 | 10 | [3] |
| *Asterocampa leilia*[a] | -29.6 | 2.7 | 1.9 | 2 | 3 | 5 | [4] |
| *Battus philenor* | 14.3 | 1.4 | 1.6 | 5 | 5 | 10 | [3] |
| *Bibio marci* | -26.3 | 7.6 | 5.6 | - | - | - | [5] |
| *Caligo eurilochus* | 5.9 | 1.7 | 1.8 | 2 | 2 | 4 | [2] |
| *Calliphora erythrocephala* | -16.4 | 2.6 | 2.1 | 1 | 1 | 2 | [6] |
| *Coenosia attenuata*[b] | 0 | 4.4 | 4.4 | 1 | 1 | 2 | [7] |
| *Colias eurytheme* | -39 | 2.4 | 1.4 | - | - | - | [8] |
| *Dilophus febrilis* | -22 | 10 | 7.8 | - | - | - | [5] |
| *Dissosteira carolina* | 24.1 | 3.8 | 4.7 | 16 | 16 | 32 | this study |
| *Operophtera brumata* | -12.5 | 6.4 | 5.6 | 10 | 10 | 20 | [9] |
| *Orgyia antiqua* | -36.6 | 6.1 | 3.8 | 8 | 6 | 14 | [10] |
| *Parthenos sylvia* | -5 | 2 | 1.9 | 2 | 2 | 4 | [2] |
| *Polygonia c-album* | -10.3 | 2.9 | 2.6 | 2 | 2 | 4 | [2] |
| *Spharagemon equale* | 30.5 | 4 | 5.2 | 15 | 15 | 30 | this study |
| *Volucella pellucens* | -22.2 | 1.8 | 1.4 | - | - | - | [11] |

a = data reported as range of individuals, midpoint used here
b = data reported visually in figure, estimated data used here

**Literature Cited for Supplemental Table 2**


1. Lau, T. F. (Stanley), Gross, E. M. & Meyer-Rochow, V. B. Sexual dimorphism and light/dark adaptation in the compound eyes of male and female Acentria ephemerella (Lepidoptera: Pyraloidea: Crambidae). *EJE* **104**, 459–470 (2007).

2. Rutowski, R. L., Gislén, L. & Warrant, E. J. Visual acuity and sensitivity increase allometrically with body size in butterflies. *Arthropod Structure & Development* **38**, 91–100 (2009).

3. Bergman, M. & Rutowski, R. L. Eye morphology and visual acuity in the pipevine swallowtail ( *Battus philenor* ) studied with a new method of measuring interommatidial angles. *Biological Journal of the Linnean Society* **117**, 646–654 (2016).

4. Rutowski, R. L. & Warrant, E. J. Visual field structure in the Empress Leilia, Asterocampa leilia (Lepidoptera, Nymphalidae): dimensions and regional variation in acuity. *Journal of Comparative Physiology A* **188**, 1–12 (2002).

5. Zeil, J. Sexual dimorphism in the visual system of flies: The compound eyes and neural superposition in bibionidae (Diptera). *Journal of comparative physiology* **150**, 379–393 (1983).

6. Land, M. F. & Eckert, H. Maps of the acute zones of fly eyes. *J. Comp. Physiol.* **156**, 525–538 (1985).

7. Gonzalez-Bellido, P. T., Wardill, T. J. & Juusola, M. Compound eyes and retinal information processing in miniature dipteran species match their specific ecological demands. *Proceedings of the National Academy of Sciences* **108**, 4224–4229 (2011).

8. Merry, J. W., Morehouse, N. I., Yturralde, K. & Rutowski, R. L. The eyes of a patrolling butterfly: Visual field and eye structure in the Orange Sulphur, Colias eurytheme (Lepidoptera, Pieridae). *Journal of Insect Physiology* **52**, 240–248 (2006).

9. Meyer-Rochow, V. B. & Lau, T. F. S. Sexual dimorphism in the compound eye of the moth Operophtera brumata (Lepidoptera, Geometridae). *Invertebrate Biology* **127**, 201–216 (2008).

10. Lau, T. F. S. & Meyer-Rochow, B. V. The compound eye of Orgyia antiqua (Lepidoptera: Lymantriidae): Sexual dimorphism and light/dark adaptational changes. *EJE* **104**, 247–258 (2007).

11. Warrant, E. J. Sensory matched filters. *Current Biology* **26**, R976–R980 (2016).
